# Macrophages Lacking TSC2 have mTORC1-dependent GPNMB Augmentation Ameliorating Cardiac Ischemia-Reperfusion Injury

**DOI:** 10.1101/2025.05.29.656917

**Authors:** Mohammad Keykhaei, Navid Koleini, Mariam Meddeb, Masih Tajdini, Malihe Rezaee, Qiao Huang, Tegbir Panesar, Mark Ranek, Luigi Adamo, David A. Kass

**Author notes:** Address Correspondence: David A. Kass, MD Division of Cardiology Ross Building Rm 858 720 Rutland Avenue Baltimore, MD 21205 @dkassjhu.

## Abstract

**Introduction:** Macrophages (MΦ) modulate both myocardial inflammatory and reparative phases following ischemia-reperfusion (I/R) injury. The mechanistic target of rapamycin (mTOR) is thought to play an important role in MΦ phenotype and functionality, but studies report conflicting net influences suggesting dependence on disease context and downstream signaling. Here, we tested the impact of MΦ with constitutive mTORC1 activation induced by targeted deletion of tuberous sclerosis complex 2 (TSC2) on cardiac responses to I/R injury.

**Methods/Results:** Myeloid TSC2 depleted (MΦ^TSC2-/-^) mice were generated by crossing Lys2^Cre^ x TSC2^flx/flx^. Bone-marrow derived MΦ^TSC2-/-^ vs control MΦ had basal increased mTORC1 and reduced mTORC2 activity. MΦ^TSC2-/-^ were differentially responsive to stimulation by lipopolysaccharide/IFN-γ or IL-4 *in vitro*, and all disparities were prevented by rapamycin confirming the model. In vivo, MΦ^TSC2-/-^ mice were strongly protected against I/R injury, with minimal change in ejection fraction, less LV dilation, hypertrophy, lung edema, or activation of stress/pro fibrotic genes. Mice pre-treated with anti-LY6G Ab to deplete neutrophils were still similarly protected, suggesting that the impact was primarily related to MΦ. MΦ^TSC2-/-^ mice had less myocardial pro-inflammatory macrophages (CCR2^+^MHC-II^hi^), LY6C^+^ monocytes, neutrophils, and CD8^+^ T cells 5 days post-I/R, and fewer CCR2^+^ but more CCR2^-^ MΦ 2 weeks post I/R. Both MΦ^TSC2-/-^ in vitro and in vivo post I/R phenotypes were converted to WT by rapamycin, supporting mTORC1 dependence. Lastly, synthesis of glycoprotein nonmetastatic melanoma protein B (GPNMB), a principally MΦ anti-inflammatory secreted protein protective against myocardial infarction was enhanced in MΦ^TSC2-/-^ macrophages and hearts following I/R in an mTORC1 dependent manner. **Conclusion:** Constitutive macrophage-specific mTORC1 activation via TSC2 deletion reduces pro-inflammatory cell infiltration, increases GPNMB protein expression and preserves heart function following I/R injury. Rapamycin eliminates these effects. These results identify a cardioprotective mTORC1-GPNMB signaling nexus in MΦ in vivo.

## Introduction

Ischemia-reperfusion (I/R) injury is a substantial contributor to heart failure worldwide, a major cause of morbidity and mortality^1^. While restoring coronary perfusion limits infarct size, it also subjects the heart to reperfusion-coupled oxidative stress, calcium overload, and pro-inflammatory signaling that further exacerbate cardiomyocyte demise and adverse remodeling^2^. The inflammatory component occurs in stages starting with acute inflammation mediated by neutrophils and C-C motif chemokine receptor 2 (CCR2+) monocytes and macrophages (MΦ) derived principally from non-resident cells. Early phase CCR2+ macrophages are generally pro-inflammatory, secreting cytokines such as tumor necrosis factor-alpha (TNF-α), interleukin-6 (IL- 6), and interleukin-1β (IL-1β) that amplify tissue remodeling and cell dissolution. Tissue-resident CCR2- MΦ favor cardiomyocyte salvage and repair ^3–6^.

A primary regulator of macrophage phenotype and function is the mechanistic target of rapamycin (mTOR), an evolutionarily conserved serine/threonine kinase complex that integrates nutrient, energy, and immune signals^7–9^. MTOR exists in two functionally distinct complexes: mTORC1 (rapamycin-sensitive), which promotes protein synthesis and cell growth via S6K1 and 4E-BP1, and mTORC2 (largely rapamycin insensitive) that controls cytoskeletal organization and metabolic adaptations notably coupled to Akt activation^10^. The tuberous sclerosis complex (TSC1/TSC2) is a major proximal constitutive suppressor of mTORC1 activity by limiting GTP-bound to the small G-protein/mTORC1 activator Rheb. Loss of function mutations in TSC1 or TSC2 underlie tuberous sclerosis syndrome, associated with constitutive mTORC1 activation that manifests with tumors, neurocognitive disease, and seizures^11^. TSC1 gene deletion results in TSC2 degradation with loss of mTORC2 signaling as well, whereas TSC2 deletion leads to negative feedback on mTORC2 but not its elimination^12, 13^.

Studies on the role of mTORC1 activity to MΦ phenotypes have revealed both pro- and anti-inflammatory behaviors. Deletion of TSC1 or TSC2 in MΦ can cause increased inflammatory modulation that worsens certain disease conditions^7, 12, 14–21^. For example, MΦ-TSC2^-/-^ mice can develop lung and cardiac sarcoidosis, MΦs contributing to granuloma formation, fibrosis, and impaired organ function in an mTORC1 dependent manner ^14, 15^. They can also evolve autoimmune arthritis (Still’s disease)^17^, and amplify pro-inflammatory cytokines and fibrosis after renal I/R injury^18^. By contrast, other studies found suppressing mTORC1 signaling by genetic mTOR or raptor gene deletion^22^ or by rapamycin^23^ is pro-inflammatory, implying mTORC1 normally confers anti-inflammatory effects. These disparate results suggest the precise manner and signaling coupled to mTOR activity in MΦ is likely context driven.

To our knowledge, no prior study has examined the role of MΦ-mTOR activation on I/R injury; however a recent report using a total coronary occlusion infarction model reported that infiltrating MΦs were cardioprotective by secreting glycoprotein non-metastatic melanoma protein B (GPNMB) ^24^. GPNMB is widely expressed in cells though most prominently by MΦs and under stressed conditions where it generally confers anti-inflammatory effects on other cells including cardiomyocytes^24, 25^. Intriguingly, GPNMB is also among the most highly upregulated genes in several TSC1/TSC2 loss of function related tumors linked to mTORC1 activation^26–28^. Whether MΦs with mTOR stimulation exhibit enhanced GPNMB production and ameliorate post I/R injury is unknown. We tested this hypothesis in mice with TSC2 selective knocked out of MΦs which indeed confer substantial cardio-protection against I/R injury coupled to enhanced GPNMB.

## Materials and Methods

### Animal Models and Generation of Macrophage-specific TSC2 Knockout Mice

MΦ^TSC2-/-^ mice were generated by crossing mice with loxP flanked TSC2 alleles (TSC2^fl/fl^) (generous gift from Michael Gambello, University of Texas Health Science Center at Houston, Houston, Texas, USA) ^29^ with mice expressing Cre recombinase under the control of the lysozyme M (Lyz2) promoter (Jackson Laboratory, Strain #004781). These mice have myeloid lineage-specific deletion of TSC2 that most prominently affects macrophages and neutrophils. Littermate TSC2^fl/fl^ mice lacking Cre expression served as controls (CON). Mice were housed in pathogen-free conditions with a 12-hour light/dark cycle with standard chow and water ad libitum. All studies were performed in compliance with the relevant ethical regulations for animal testing and research and the protocols approved by the Johns Hopkins School of Medicine Committee on Animal Care (IACUC). Male mice of age 8-10 weeks were used in the study.

### In Vivo Ischemia-Reperfusion (IR) Model

I/R was generated using a minimal surgical approach ^30^ with mice anesthetized with 3% isoflurane inhalation but not ventilated, the heart temporarily externalized via a small hole in the 4th intercostal space, and subjected to reversible ligation (slip-knot) of the LAD with a 6-0 suture. The heart was immediately returned to the mediastinum, air evacuated and skin closed. After 45- 60 min of ischemia, the heart was re-externalized, reperfusion obtained by releasing the slip knot, and then rapidly returned, air removed, and the chest incision closed. Animals were monitored and received one dose of buprenorphine (0.03mg/kg) for pain. Beyond the ischemic time, the procedure took ∼5 min with mortality <5%.

The I/R protocol was performed in three different cohorts: 1) control and MΦ^TSC2-/-^ mice; 2) the same groups pre-administered 400 μg anti-Ly-6G antibody (clone 1A8, BioXCell) by i.p. injection initiated 24 hours prior to I/R and dosed 2x/week for 4 weeks (neutrophil depletion group); and 3) control and MΦ^TSC2-/-^ randomized to receive vehicle (3.3 mL/kg of 0.9% NaCl, 5% PEG400, 5% Tween 20, and 3% ethanol) or rapamycin (Target Mol Cat#53123-88-9, 2 mg/kg in vehicle) given daily by intraperitoneal injection. Treatment started the day of I/R and continued for 4 weeks.

### Echocardiography Analysis

M-mode transthoracic echocardiography was performed in conscious mice using a VisualSonics Vevo 2100 system with an 18-38mHz transducer (SanoSite Inc) by an operator blinded to study conditions. Left ventricular diameter at end-diastole and end-systole (LVIDD, LVESD), ejection fraction (EF), and average wall thickness were determined using VisualSonics software in a blinded manner. Values represent averages from five or more consecutive heartbeats.

### Blood Pressure Measurement

Systolic blood pressure in conscious mice was measured by tail-cuff plethysmography (CODA, Kent Scientific) following manufacturer’s protocol. Data from last 5 of 10 independent measures/mouse are reported, at ambient temperature at 31°C to ensure proper tail blood flow.

### Bone Marrow-Derived Macrophages (BMDM) Isolation and Culture

BMDMs were isolated from mice following CO₂ euthanasia. Femurs and tibias were aseptically removed, and bone marrow cells flushed out with PBS delivered by sterile 25G needle. Cells were passed through a 70-μm strainer to remove debris and centrifuged at 500g for 5 minutes at 4°C. Cells were then resuspended in RPMI 1640 medium (Thermo Fisher Cat# 11875119) supplemented with 10% fetal bovine serum (Sigma-Aldrich Cat# F0392), 1% penicillin/streptomycin (Thermo Fisher Cat# 15140122), and 2 mM L-glutamine (Thermo Fisher Cat# 25030081). Cells were plated in bacteriological-grade Petri dishes and cultured in the presence of macrophage colony-stimulating factor (M-CSF) (PeproTech) for 7 days. The medium was replaced on days 3 and 5 to ensure optimal macrophage differentiation. On day 7, adherent macrophages were harvested in cold PBS with gentle scraping and plated for subsequent study. Cells were cultured at 37 °C in a humidified CO2 (5%) incubator. To induce a pro-inflammatory response, cells received 10 ng/mL lipopolysaccharide (LPS, Invitrogen) and 20 ng/mL interferon gamma (IFN-γ, Invitrogen) for 24 hours. To induce an anti-inflammatory response, cells received 40 ng/mL IL-4 (PeproTech) for 24 hours. In some studies, cells were additionally incubated with rapamycin (Calbiochem) added to the medium (final concentration 100 nM).

### Macrophage Proliferation and Morphometric Analysis

Macrophage proliferation was measured in BMDMs cultured for 10 days by counting the total number of adherent cells per well. Cell area was quantified with ImageJ (Fiji) software, using phase-contrast images acquired at 10× magnification, with at least 50 cells measured per condition.

### Protein Expression Analysis

BMDMs were collected in SDS lysis buffer supplemented with a 1:100 dilution of Halt Protease and Phosphatase Inhibitor Cocktail (100X) (Thermo Scientific, #78446). Lysates were sonicated, heated, and centrifuged at 14,000g for 15 minutes to remove cellular debris. Protein concentration in the supernatants was measured using the bicinchoninic acid (BCA) assay. Sodium dodecyl sulfate–polyacrylamide gel electrophoresis was used for protein separation using 4-15% Criterion TGX Gel 18W 30ul (Bio-Rad Cat# 5671084) or 4-20% Criterion TGX Gel 26W 15ul (Bio-Rad Cat# 5671095). Proteins were blotted onto nitrocellulose membrane 0.45 µm, 8.5 x 13.5 cm (Bio-Rad Cat#1620167) using a semi-dry Trans-Blot® Turbo™ Transfer System (BioRad Cat#1704150). Antibodies from Cell Signaling were used for P-P70S6 Kinase (9205, 1/000), P70S6 Kinase (9202, 1/1000), TSC2 (4308, 1/1000), P-4EBP1 (9451, 1/1000), 4EBP1 (9644, 1/1000), P-AKT (Ser473) (4060, 1/1000), P-AKT (Thr308) (5106, 1/1000), AKT (9272, 1/1000), and GPNMB (90205, 1/200). To prevent nonspecific binding, membranes were blocked for 1 hour at room temperature with Intercept® (TBS) Blocking Buffer (Li-COR) followed by extensive washing in TBS-Tween (TBS-T). Membranes were incubated overnight at 4°C with the primary antibody diluted 1:1000 in TBS antibody diluent (Li-COR), followed by washing in TBS- T the next morning. Membranes were then incubated for 1 hour at room temperature with secondary antibody (Goat anti-mouse and rabbit secondary antibodies, Li-COR: Cat# 926-32210, Cat# 926-32211, Cat# 926-68071, Cat# 926-68070) at a 1:5000 dilution. Membranes underwent 1-hour wash in TBS-T before imaging. Protein bands were visualized using a Li-Cor Odyssey imager, and densitometry analysis was conducted with Image J (Version 1.53). Densitometry data were normalized first to total protein in each gel and subsequently to the non-failing control group, which was assigned a mean value of 1.0. Total protein concentration was determined using the Revert 520 Total Protein Stain Kit (Li-COR) according to the manufacturer’s instructions.

### RNA Extraction and Semiquantitative Real Time RT-PCR

Total RNA was extracted with TRIzol® reagent (Thermo Fisher Scientific) following the manufacturer’s protocol. 100 ng of RNA was reverse transcribed into complementary DNA (cDNA) using the High-Capacity cDNA Reverse Transcription Kit (Applied Biosystems, #4368814). Quantitative real-time PCR (qRT-PCR) was conducted on a QuantStudio 5 system (Thermo Fisher Scientific) utilizing the TaqMan Fast Advanced Master Mix (Applied Biosystems, #4444963) and specific TaqMan primers (detailed in **Supplementary. Table1**). Gene expression levels were normalized to the housekeeping gene RPS18, and relative expression changes were analyzed using the 2^−ΔΔCt^ method, with experimental conditions compared to controls.

### GPNMB ELISA assay

Cytokine levels of IL-10 and TNF-α, were quantified with ELISA kits (BioLegend) GPNMB was measured using ELISA kit (Thermo Fisher, Cat# EM59RB). Experimental supernatants were collected and centrifuged at 3,000g for 5 minutes to remove debris. The clarified supernatants were analyzed in duplicate following the manufacturer’s protocol.

### Histological Analysis

Four weeks after I/R, mice were euthanized, hearts perfused with PBS to remove residual blood, then fixed overnight at 4°C in 4% paraformaldehyde (PFA) in PBS, followed by dehydration in 70% ethanol before paraffin embedding. Transverse LV sections were cut into 4- µm thick slices and mounted on positively charged glass slides. Masson’s Trichrome stain (Richard-Allan Scientific, #87020) was used to assess for fibrosis, with whole heart sections imaged and analyzed using ImageJ (Fiji).

### Flow Cytometry and Cell Sorting

Hearts were perfused with PBS, weighed, minced, and enzymatically digested at 37°C for 45 minutes in DMEM (Gibco, 11965-084) containing Collagenase IV (4500 U/ml, Millipore Sigma C5138), Hyaluronidase I (2400 U/ml, Millipore Sigma H3506), and DNAse I (6000 U/ml, Millipore Sigma D4527). The digestion reaction was stopped using HBSS supplemented with 2% FBS and 0.2% BSA, followed by filtration through 40-µm strainers. Red blood cells were lysed by incubating the cell suspension with ACK lysis buffer (Gibco A10492-01) for 5 minutes at room temperature, after which cells were washed with DMEM and resuspended in 100 µl of FACS buffer (PBS containing 2% FBS and 2 mM EDTA). For immunostaining, cells were incubated for 30 minutes at 4°C with fluorescence-conjugated monoclonal antibodies at 1:200 dilution, followed by washing and resuspension in 250 µl of FACS buffer for analysis. All experiments were performed on a Cytek® Aurora Full Spectrum Flow Cytometric Analyzer (JHU Ross Flow cytometry core) and analyzed using FlowJo software analysis. For monocyte, neutrophil, macrophage, and lymphocytes analysis, antibodies used included: PerCP/Cyanine5.5 anti-mouse CD45 Antibody (BioLegened, 103132), PE/Cyanine7 anti-mouse CD64 (FcγRI) Antibody (BioLegened, 139314), APC/Cyanine7 anti-mouse Ly-6G Antibody (BioLegened, 127624), Alexa Fluor® 700 anti-mouse/human CD11b Antibody (BioLegened, 101222), Brilliant Violet 650™ anti-mouse Ly-6C Antibody (BioLegened, 128049), APC anti-mouse CD192 (CCR2) Recombinant Antibody (BioLegened, 160103), Brilliant Violet 711™ anti-mouse I-A/I-E Antibody (BioLegened, 107643), Brilliant Violet 785™ anti-mouse CD8a Antibody (BioLegened, 100749), Spark NIR™ 685 anti-mouse CD4 Antibody (BioLegened, 100476), Brilliant Violet 421™ anti-mouse CD19 Antibody (BioLegened, 152421).

### Statistical Analysis

Data are presented as mean ±SEM. Statistical analyses were performed with Prism (GraphPad, Version 10), with tests and sample sizes for each group detailed in the figures or figure legends. For two-way comparisons, Student’s two-tailed t-test was used, for multiple group comparisons, one- or two-way analysis of variance (ANOVA) with appropriate multiple comparisons test used. For non-normally distributed data, a Kruskal-Wallis test was used. Exact p-values are provided in figures and/or legends as well as statistical tests.

## Results

### Characterization of MΦ^TSC2-/-^ BMDM in vitro

The MΦ^TSC2-/-^ model (**Supplementary** Fig 1a) generated BMDMs that essentially lacked TSC2 (**Fig. 1a**), that had constitutive mTORC1 activation reflected by greater phosphorylation of downstream proteins p70S6K1 and 4E-BP1 (**Fig. 1b, Supplemental Fig. 1b**). Corresponding mTORC2 activity was reduced, as reflected by reduced Akt phosphorylation at S473 and T308 (**Fig. 1c, Supplemental Fig. 1c**). Both changes were eliminated with rapamycin incubation, confirming the role of mTORC1 activation and negative feedback of mTORC1 on mTORC2.

**Figure 1.**
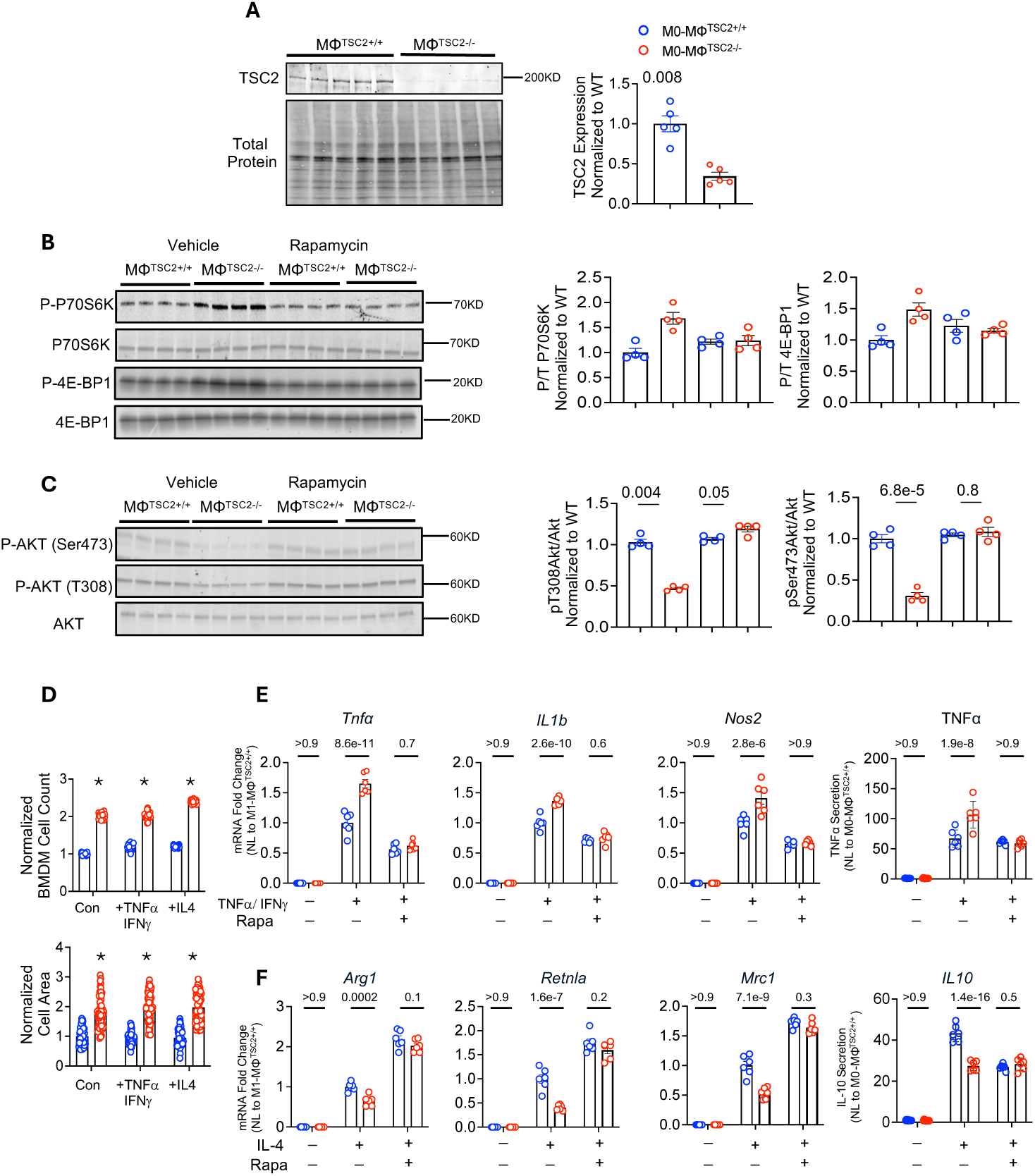
TSC2 deletion in macrophages causes constitutive mTORC1 activation altering in vitro macrophage stimulation responses **A**, Immunoblot of TSC2 protein expression in BMDMs from MΦTSC2^+/+^versus MΦTSC2^-/-^ (n=5 per group, unpaired t-test). **B**, *Left:* Immunoblot for P70S6K and 4E-BP1 phosphorylation and total protein in BMDMs from same two models incubated with vehicle or rapamycin. (n=4 per group); *Right:* phospho/total protein. **C**, *Left:* Immunoblot of Akt activation from same experiment (n=4 per group), *Right:* Summary data; all analyzed by one-way ANOVA, Dunnett multiple correction test. **D**, Proliferation of BMDMs for both groups without vehicle, LPS/IFN-γ, or IL-4 stimulation. *Upper:* cell numbers (n=20/group), *Lower:* cell size (n=60/group). For all, data normalized to MΦTSC2^+/+^ results for each stimulus, two-way ANOVA with Šidák multiple comparisons test. **E**, Response of BMDMs from both models to IFN-γ+ LPS stimulation ± rapamycin (n=6/group), **F** to IL-4 ± rapamycin (n=6/group), two-way ANOVA with Šidák multiple comparison test. Data are mean±SEM. *P<0.05. 4E-BP1, eukaryotic translation initiation factor 4E-binding protein 1; P70S6K, ribosomal protein S6 kinase; Arg1, arginase 1; LPS, lipopolysaccharide; Mrc1, mannose receptor C-type 1; Nos2, nitric oxide synthase 2; Retnla, resistin-like alpha; TNFα, tumor necrosis factor alpha, other abbreviations in text.

BMDM proliferative growth was measured in vitro while incubated with vehicle control, LPS/IFN-γ, or IL-4. MΦ^TSC2-/-^ BMDMs were more proliferative and had larger cell area with each condition versus MΦ^TSC2+/+^ controls (**Fig. 1d**). LPS/ IFN-γ stimulated MΦ^TSC2-/-^ BMDMs stimulated with LPS/IFN-γ had higher expression of *Tnfa*, *Nos2*, and *Il1b* and TNF-α secretion (**Fig. 1e**), whereas IL-4 stimulated MΦ^TSC2-/-^ BMDMs had less expression of *Arg1*, *Retnla*, and *Mrc1* and IL-10 secretion (**Fig. 1f**). These differences were also abrogated by co-treatment with rapamycin, supporting their mTORC1-dependence. These results support the model as designed.

### MΦ^TSC2-/-^ hearts exhibit improved function, lung edema, and stress responses post I/R

Echocardiography measured 4-weeks post I/R revealed substantial cardio-protection in MΦ^TSC2-/-^ versus CON mice, with EF declining from 81% to 50% in WT yet barely changing in MΦ^TSC2-/-^. The latter was related to a lack of end-diastolic chamber dilation and less incrased in end-systole volume in MΦ^TSC2-/-^ post I/R (**Fig. 2a, b**). Wall thickness **(Fig. 2b)**, systolic blood pressure and myocardial mass (**Supplemental Fig. 1d**) were unchanged in either model. Wet/dry lung weight, an indicator of pulmonary edema rose in the control but not MΦ^TSC2-/-^ mice (**Fig. 2c**).

**Figure 2.**
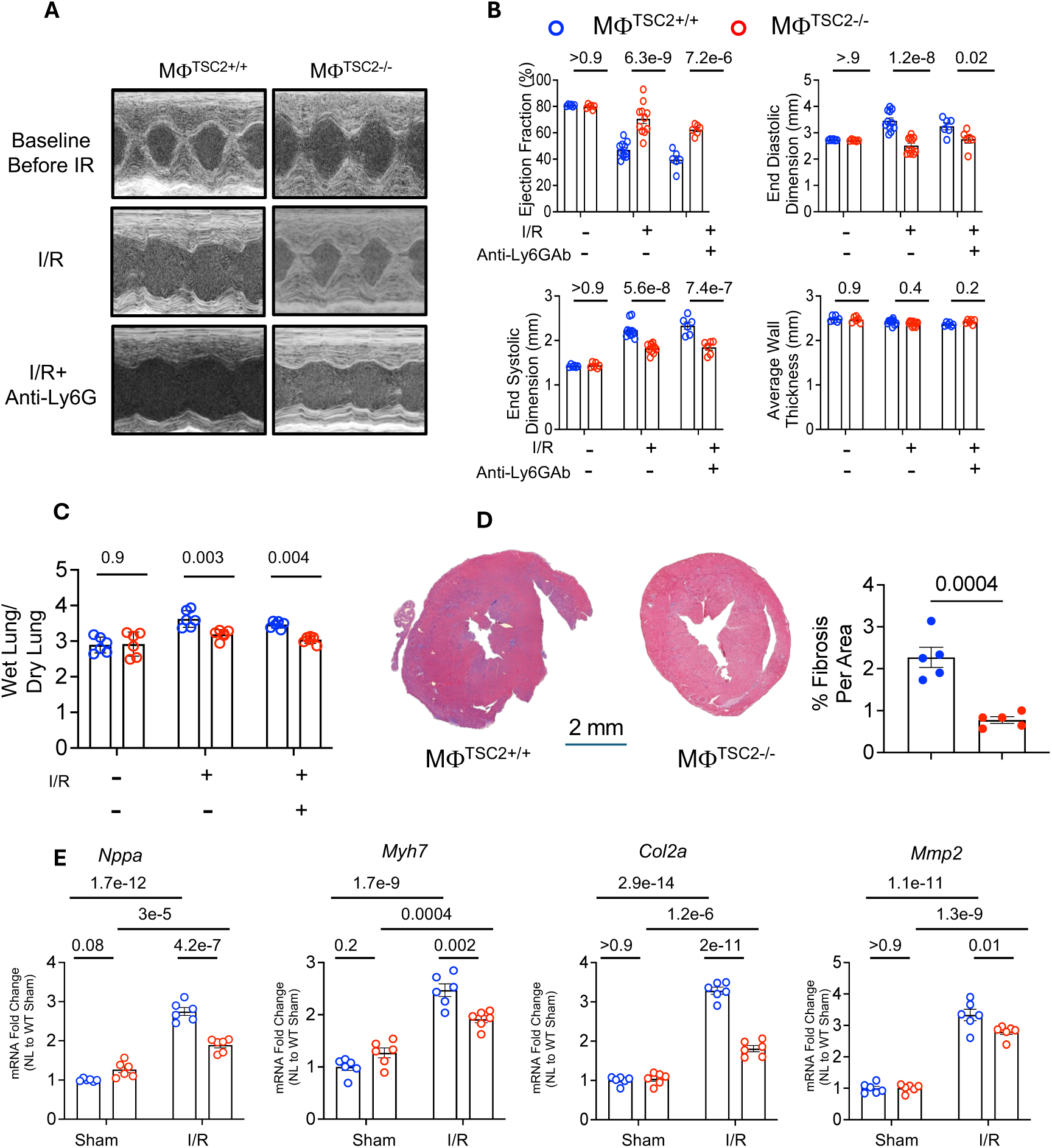
MΦTSC2^-/-^ mice are protected against post-I/R cardiac dysfunction **A**, Representative echocardiographic images at baseline and 4 weeks post– I/R with or without anti-Ly6G antibody pretreatment in MΦTSC2^+/+^ and MΦTSC2^-/-^ mice. **B**, Quantification of left ventricular function, cross sectional dimensions, and wall thickness (n=6 per group). **C**, Lung wet-to-dry weight ratios (n=6 per group). **D**, Masson’s trichrome stained cross section with percent fibrosit area (n=5 per group, unpaired t-test). **E**, Expression of hypertrophic and matrix remodeling genes (A-type natriuretic peptide, Nppa, β-myosin heavy chain, Myh7, collagen type 2a1, Col2a1, metalloproteinase 2, Mmp2) in MΦTSC2^-/-^ hearts vs controls with either Sham or I/R (n=6 per group. Data are shown as mean ± SEM. Col2a1, collagen type II alpha 1 chain; EF, ejection fraction; I/R, ischemia–reperfusion; Ly6G, lymphocyte antigen 6 complex locus G; MΦ, macrophage; Mmp2, matrix metallopeptidase 2; Myh7, myosin heavy chain 7; Nppa, natriuretic peptide A; SEM, standard error of the mean; TSC2, tuberous sclerosis complex 2. Panels B, C, and E analysis using two-way ANOVA with Šidák multiple comparisons test.

LysM-Cre is prominently expressed in neutrophils and macrophages. To determine if the cardioprotection involved TSC2-KO in neutrophils, experiments were repeated in mice pre-treated with anti-Ly6G antibody to first deplete neutrophils. These results are also displayed in **Fig. 2a, b**. There was similar cardio-protection against I/R in neutrophil-depleted MΦ^TSC2-/-^ mice.

Histopathology of post I/R hearts revealed fairly little fibrosis, but it was still significantly more in CON versus MΦ^TSC2-/-^ (**Fig. 2d**). Gene markers of cardiac stress (*Nppa, Myh7)* and fibrosis/matrix remodeling (*Col2a1*, and *Mmp2*) also rose more in CON versus MΦ^TSC2-/-^ (**Fig. 2e**).

### MΦ^TSC2-/-^ mice have reduced proinflammatory immune cell infiltration following I/R

Myocardial monocyte infiltration typically peaks ∼5 days after IR injury^31^ so we assessed monocyte and other inflammatory cell recruitment in CON and MΦ^TSC2-/-^ mice at this time. The total number of immune cells (CD45+) was comparable between MΦ^TSC2-/-^ and controls (**Fig. 3a**); however, MΦ (CD45^+^Ly6G^-^CD11b^+^CD64^+^) were significantly reduced in MΦ^TSC2-/-^ hearts (**Fig. 3b**). There were also less inflammatory monocytes (Ly6C^+^) and macrophages (CCR2+) in MΦ^TSC2-/-^ (**Fig. 3c, d**). The more pro-inflammatory infiltrating CCR2+ MHCII^hi^ macrophages were reduced in relative number in MΦ^TSC2-/-^ mice vs CON, while resident more reparative CCR2- MHCII^low^ cells were increased (**Fig. 3e**). Two weeks post-IR, overall leukocyte (CD45+) and macrophage (CD64+) infiltration was comparable between groups. However, in MΦ^TSC2⁻/⁻^ hearts, there were less pro-inflammatory CCR2⁺ MHCII^hi^ macrophages **(Supplementary** Fig. 2a-c). We confirmed no differences in immune cell phenotypes for both groups subjected to sham procedure (**Supplementary** Fig. 2d-f).

**Figure 3.**
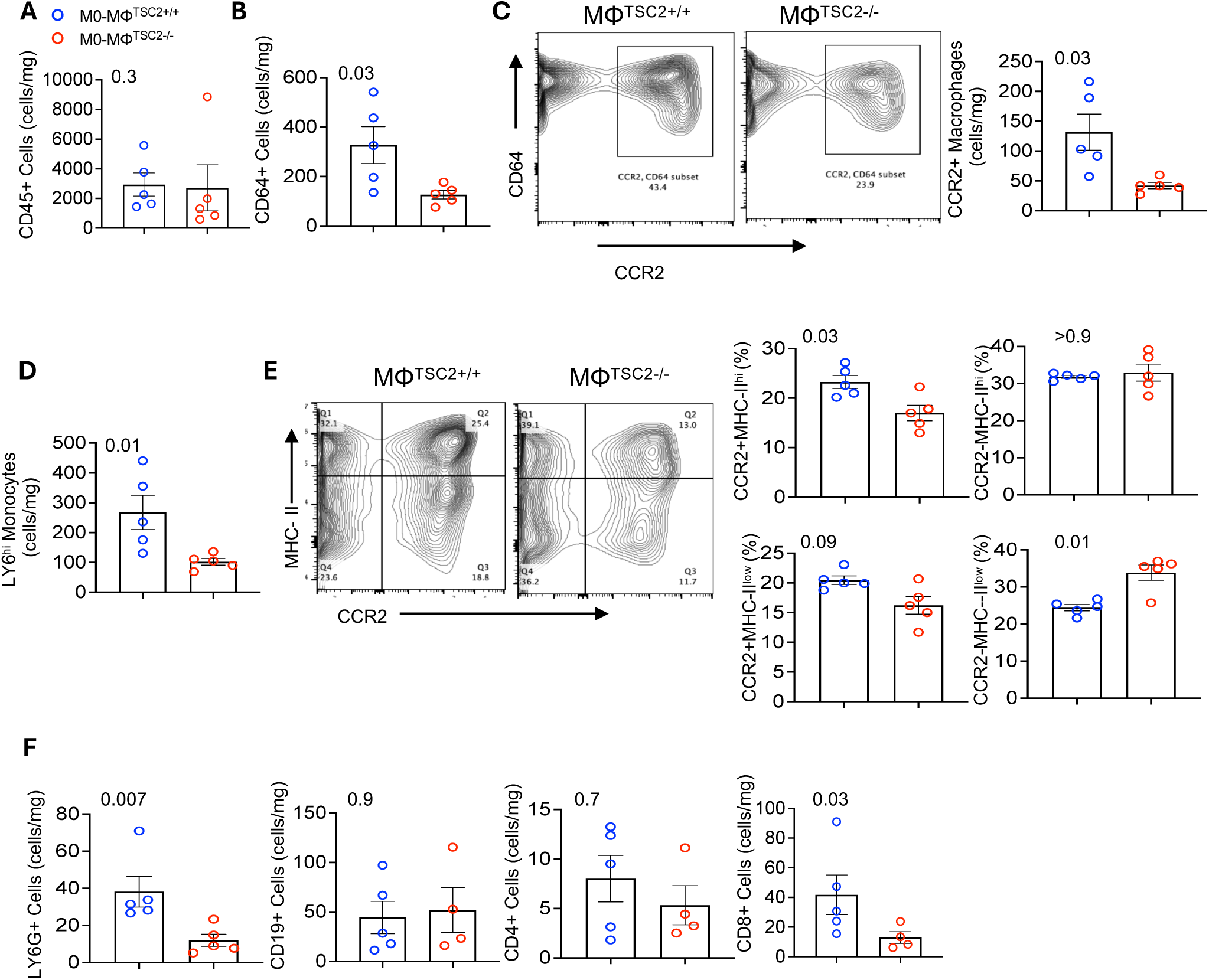
MΦTSC2^⁻/⁻^ mice have reduced pro-inflammatory immune cell infiltration in the heart 5 days after ischemia-reperfusion. **A**, Number of CD45⁺ per mg tissue between MΦTSC2⁺/⁺ and MΦTSC2⁻/⁻ hearts (n=5/group). **B**, Number of CD64⁺ macrophages per mg in hearts from both models (n=5/group). **C**, Representative flow cytometry density plots for CCR2⁺ / CD64⁺ macrophages (n=5/group). **D**, Number of Ly6C⁺ monocytes per mg tissue in both groups, (n=5/group). **E**, Representative flow cytometry analysis of macrophage subsets (n=5/group). **F**, Ly6G⁺ (neutrophils), CD8⁺ and CD4^+^ (T cells), and CD19⁺ (B cells) each normalized to mg tissue for both groups (n=5). Analysis used two-tailed unpaired t-tests. Data are mean ± SEM. CD45, leukocyte common antigen; CCR2, C-C chemokine receptor type 2; CD64, Fc gamma receptor 1; Ly6C, lymphocyte antigen 6 complex locus C; Ly6G, lymphocyte antigen 6 complex locus G; MHCII, major histocompatibility complex class II; SEM, standard error of the mean; TSC2, tuberous sclerosis complex 2.

CD4+ T-cells (CD45^+^CD4^+^) and B-cells (CD45^+^CD19^+^) infiltration did not differ significantly between groups 5 days post I/R, but neutrophil infiltration (CD45^+^Ly6^+^) and CD8^+^ T-cell infiltration (CD45^+^CD3^+^CD8^+^) were significantly reduced in MΦ^TSC2-/-^ mice (**Fig. 3f)**.

### Rapamycin blocks protective effects of MΦ^TSC2-/-^ on cardiac I/R injury

To test the requirement of mTORC1 to the cardio-protection and altered immune landscape post I/R, studies were repeated in the presence of rapamycin. Echocardiographic data no longer showed any difference between MΦ^TSC2-/-^ mice and controls (**Fig. 4a, b**), and lung wet/dry weight ratio was similarly elevated in both groups (**Fig. 4c**) as was myocardial fibrosis (**Fig. 4d**) and cardiac stress and fibrosis/matrix gene expression (**Fig. 4e**). Systolic blood pressure and myocardial mass remained similar between groups (**Supplemental Fig. 3**). Thus, the protective influence in of MΦ^TSC2-/-^ mice required activation of mTORC1.

**Figure 4.**
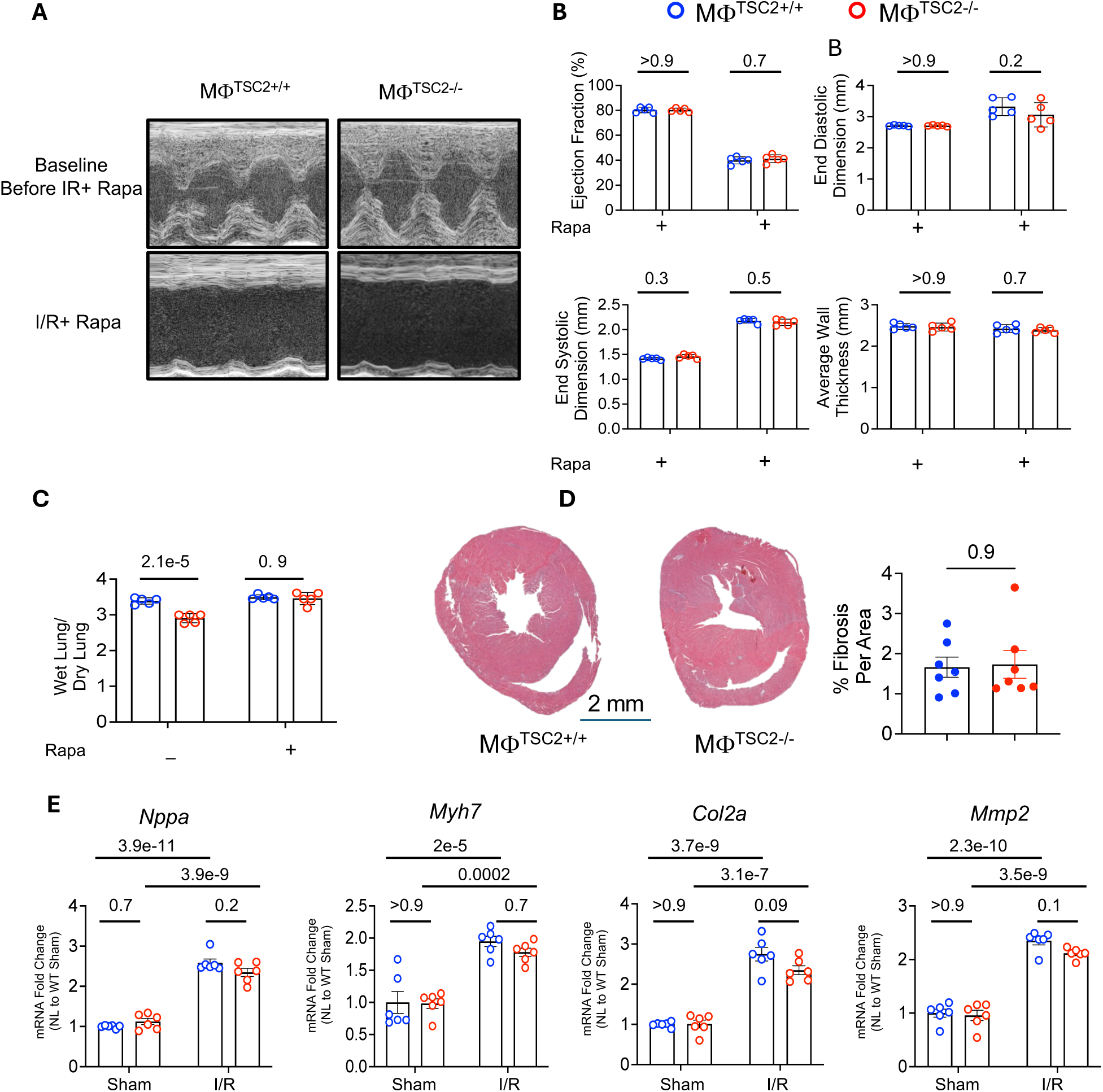
Rapamycin prevents protection against cardiac I/R injury in MΦTSC2⁻/⁻ mice. **A, B**, Example M-mode echocardiograms and summary data at 4 weeks post sham or I/R intervention in MΦTSC2⁺/⁺ and MΦTSC2⁻/⁻ mice treated with either vehicle or rapamycin (n=5 per group). Same parameters and analysis as in Figure 2. **C**, Lung wet-to-dry weight ratio, (n=5). **D**, Representative heart sections and fibrotic area quantitation (n=7, unpaired t-test). **E**, Gene expression of hypertrophy and matrix remodeling genes in same hearts (n=6, two-way ANOVA with Šidák multiple comparison test.

### MΦ^TSC2-/-^ have mTORC1-dependent increased GPNMB synthesis

In 2025, Ramadoss et al^24^ reported MΦ synthesis of GPNMB is an important mechanism of protection against myocardial infarction. In other studies primarily of neuronal, renal, and pulmonary cancers due to TSC1 deletion, GPNMB expression was among the most upregulated genes found in RNAseq analysis, and this required increased mTORC1 activity ^26, 28, 32–34^. This led us to hypothesize that GPNMB may be augmented in MΦ^TSC2-/-^ as well to enhance cardiac recovery post I/R in a rapamycin-dependent manner. In BMDMs from MΦ^TSC2-/-^ mice versus controls, GPNMB protein was increased, and this difference was absent after rapamycin treatment (**Fig. 5a, b**). In post I/R myocardium, GPNMB was also increased in MΦ^TSC2-/-^ but not CON hearts and this too was prevented by rapamycin. (**Fig. 5c**). The in vivo results were further confirmed by ELISA assay (**Fig. 5d**). Together, these findings support enhanced GPNMB secretion in resting and in vivo MΦ^TSC2-/-^ in a MTORC1-dependent manner, coupled to post I/R cardio-protection.

**Figure 5.**
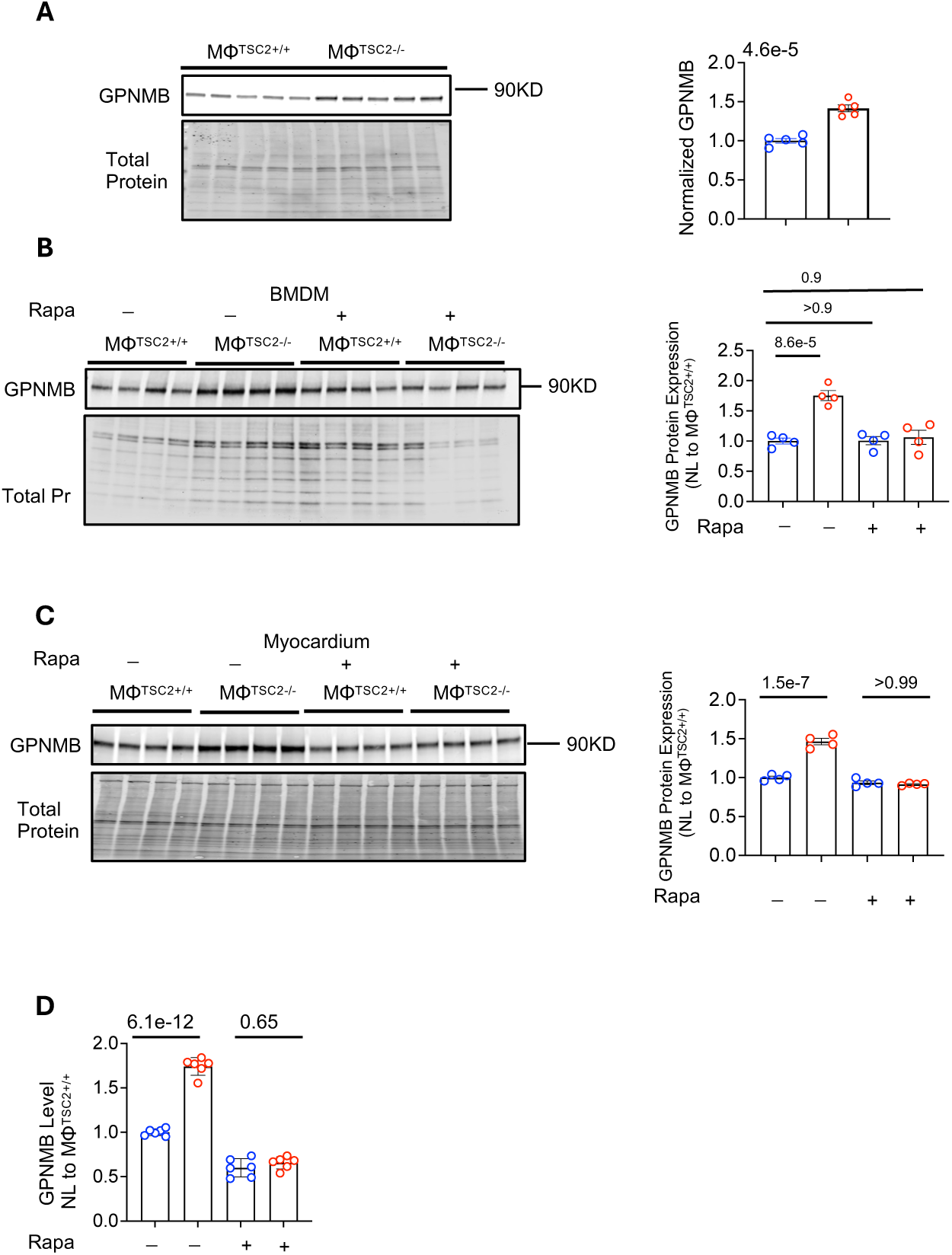
MΦTSC2⁻/⁻ mice have mTORC1-dependent increased GPNMB synthesis. **A**, Western blot and quantification show increased GPNMB protein levels in BMDMs from MΦTSC2⁻/⁻ mice compared to MΦTSC2⁺/⁺ controls under basal conditions (n=5 per group). Statistical significance was determined using unpaired two-tailed t-test. **B**, GPNMB levels remained elevated in MΦTSC2⁻/⁻ BMDMs but were significantly reduced by rapamycin treatment (n=4 per group). One-way ANOVA was used with Dunnett correction for multiple comparisons. **C**, Western blot of cardiac tissue lysates 4 weeks post–I/R revealed increased GPNMB expression in MΦTSC2⁻/⁻ hearts, which was abolished by rapamycin treatment (n=4 per group). One-way ANOVA with Dunnett correction. **D**, ELISA quantification of GPNMB protein from the same cardiac tissues confirmed the elevation in MΦTSC2⁻/⁻ hearts and its suppression by rapamycin (n=6 per group). One-way ANOVA with Dunnett correction. Data are presented as mean ± SEM. BMDM, bone marrow-derived macrophage; CON, control; ELISA, enzyme-linked immunosorbent assay; GPNMB, glycoprotein non-metastatic melanoma protein B; I/R, ischemia-reperfusion; MΦ, macrophage; mTORC1, mechanistic target of rapamycin complex 1; Rapa, rapamycin; SEM, standard error of the mean; TSC2, tuberous sclerosis complex 2.

## DISCUSSION

This study shows macrophage-specific deletion of TSC2 reduces post I/R myocardial inflammatory cell counts and corresponding LV dysfunction and remodeling in vivo. Overall monocyte numbers and in particular CCR2^+^ and CCR2^+^MHC-II^hi^ subsets, CD8^+^ lymphocytes, and neutrophils were all reduced in MΦ^TSC2-/-^ hearts after I/R, and this as well as cardiac functional benefit was prevented by rapamycin supporting the link to MΦ mTORC1 activation. GPNMB was secreted at higher levels in BMD MΦ^TSC2-/-^ and in post-I/R MΦ^TSC2-/-^ myocardium from MΦ^TSC2-/-^ mice, and both were prevented by rapamycin.

These findings reveal a complex influence of macrophage TSC2 deletion and mTORC1 activation, with pro-inflammatory bias *in vitro* yet anti-inflammatory effects *in vivo* in the setting of myocardial I/R. Prior studies investigating macrophage-specific deletion of TSC1 or TSC2 found various though generally pro-inflammatory responses in vivo ^7, 12, 14–21^ Macrophage-specific TSC2 deletion induced pulmonary and cardiac sarcoidosis, with macrophage hypertrophy, granuloma formation, and enhanced glycolysis-dependent proliferation via mTORC1-mediated CDK4 activation ^14^ ^15^. Transcriptional profiling of human cardiac sarcoidosis intriguingly reported GPNMD as a biomarker for multinucleated giant cells, coupled to mTOR activation^35^. Other studies of TSC2^-/-^ MΦ have found worsen aortic aneurysm and polyarthritis linked to inflammatory cytokines and matrix metalloproteinase activation driven by mTORC1 ^16, 17^. Such data would seem to predict a worse outcome in MΦ^TSC2-/-^ hearts after I/R. However, there are increasing data showing that the in vitro macrophage phenotype and response to cytokines poorly predict in vivo responses as the latter integrate complex signaling in the tissue and are condition specific^6^.

Prior studies involving TSC1 deletion to activate mTORC1 have also observed in vitro pro-inflammatory phenotypes but in vivo immune suppression by various other mechanisms. For example, TSC1^-/-^ macrophages protected against allergic asthma by disrupting alternative macrophage activation^19^. In another study, MΦ^TSC1-/-^ increased acute-phase inflammation and cytokine production in a renal ischemia-reperfusion injury model but reduced chronic-phase fibrosis^18^. Along with their pro-inflammatory in vitro phenotype, MΦ^TSC1-/-^ have been found to impede macrophage migration and increase apoptosis, which could dominate a net reduction of inflammation ^21^. Such changes could well have occurred in the current study as more resident versus BMD MΦ (e.g. CCR2^+^ and in particular CCR2^+^MHC-II^hi^) were found in the myocardium following I/R. These MΦ are classically pro-inflammatory^36, 37^, and CCR2 blockade or genetic deletion has been shown to improve cardiac outcome by mitigating inflammatory infiltration^38, 39^. Our finding that this requires mTORC1 activation is consistent with a prior study where this activity was suppressed by genetic deletion of mTOR or the mTORC1 complex protein Raptor^22^. Both in vitro and in vivo responses to LPS or viral infection were worse, indicating anti-inflammatory effects of mTORC1 activation. Rapamycin is also reported to be pro-inflammatory by blocking mTORC1-dependent anti-inflammatory effects of glucocorticoids on myeloid cells ^23^. GPNMB is expressed in multiple cell types but most abundantly in MΦs and glial cells, but levels rise particularly in MΦs in the setting of inflammation or tissue repair. It is a multifunctional glycoprotein that signals by Akt and Erk MAP kinase and NF-kB cascades^27^, and while augmented in inflammatory syndromes, is generally thought to confer anti-inflammatory effects ^24, 25, 32^. In tumor biology, elevated GPNMB facilitates immune evasion and contributes to check-point inhibitor resistance by modulating macrophage polarization towards a more immunosuppressive phenotype ^40^. Similarly, in lymphangioleiomyomatosis, enhanced GPNMB expression underlies disease pathogenesis through direct interactions with mTOR signaling pathways, promoting aberrant cell proliferation and tissue remodeling ^32^. GPNMB was recently reported to play a role in myocardial infarction, where its generation and release from bone-marrow derived macrophages facilitated scar formation, reduced inflammation, and helped clear apoptotic cardiomyocytes^24^. While previously shown to engage CD44-dependent signaling^41^, this study^24^ found GPNMB engaged an orphan G-protein coupled receptor GPR39 as contributing to its downstream efficacy in cardiomyocytes. However, this was not the sole mechanism, as benefits of GPNMB persisted in GPR39 KO hearts. A link between mTORC1 activation and GPNMB secretion has been suggested in studies of sarcoidosis^15^, and tumors associated with TSC1 deletion including astrocytoma^33^ and renal tumors^28^. However, the precise signaling mechanism linking the two remains unknown. Thus, our study supports a role of increased mTORC1-dependent GPNMB release to myocardial protection and anti-inflammation by MΦ^TSC2-/-^, the precise mechanism remains to be determined by future studies. Our results do raise caution when applying rapamycin in an I/R setting, as it could blunt GPNMB levels and have a potential pro-inflammatory impact.

While this study identified enhanced GPNMB and I/R cardioprotection from MΦ^TSC2-/-^ requires mTORC1 activation, it does not prove protection and anti-inflammatory changes observed absolutely requires GPNMB. We considered crossing MΦ^TSC2-/-^ with GPNMB^-/-^ mice or using adoptive cell transfer after bone marrow ablation with macrophages lacking both TSC2 and GPNMB. However, as MΦ^GPNBM-/-^ have already been shown to worsen myocardial infarction^24^ so such studies would not be definitive. Ideally, if one knew just how mTORC1 activation stimulates GPNMB synthesis, and this could be genetically prevented, then a more specific test of its role in the current setting could be performed. This remains for future studies. Lastly, LyzM is also expressed in sub-groups of dendritic cells; however, unlike macrophages and neutrophils that play major roles after I/R injury, little is known about the role of dendritic cells.

In conclusion, we show MΦ^TSC2-/-^ with stimulated mTORC1-GPNMB signaling suppresses myocardial inflammation, immune cell infiltration, and enhances cardiac functional recovery following I/R injury. This occurs despite in vitro phenotyping of these cells showing a balance in favor of greater pro-inflammatory and less anti-inflammatory activation. The results further reveal that rapamycin, conventionally viewed as an immune suppressant can have pro-inflammatory effects depending on which cell types mTORC1 activity was enhanced and its impact.

## Acknowledgments

Cytek® Aurora is funded by a NIH Grant (S10OD026859) and was used at the Johns Hopkins University Ross Flow Cytometry Core.

## Supplemental Materials

Supplemental Figures S1-S3 Table S1

## Non-standard Abbreviations and Acronyms

AKT: protein kinase B
Arg1: arginase 1
BCA: bicinchoninic acid
BMDM: bone marrow–derived macrophage
BSA: bovine serum albumin
CD11b: cluster of differentiation 11b
CD19: cluster of differentiation 19
CD4: cluster of differentiation 4
CD45: cluster of differentiation 45 (leukocyte common antigen)
CD64: Fc gamma receptor I (high-affinity IgG receptor)
CD8: cluster of differentiation 8
Col2a1: collagen type II alpha 1 chain CON control
CCR2: C-C motif chemokine receptor 2
CXCL9: C-X-C motif chemokine ligand 9
DMEM: Dulbecco’s modified Eagle medium
EDTA: ethylenediaminetetraacetic acid
EF: ejection fraction
ELISA: enzyme-linked immunosorbent assay
FACS: fluorescence-activated cell sorting
FBS: fetal bovine serum
GPNMB: glycoprotein non-metastatic melanoma protein
B HBSS: Hank’s balanced salt solution
IFN-γ: interferon gamma
IL-1β: interleukin-1 beta
IL-4: interleukin-4
IL-6: interleukin-6
IL-10: interleukin-10
I/R: ischemia-reperfusion
LPS: lipopolysaccharide
LV: left ventricle
Ly6C: lymphocyte antigen 6 complex, locus
C Ly6G: lymphocyte antigen 6 complex, locus
G MHCII: major histocompatibility complex class
II Mmp2: matrix metallopeptidase 2
MΦ: macrophage
MΦTSC2-/-: macrophage-specific
TSC2: knockout
MRC1: mannose receptor C-type 1
mTOR: mechanistic target of rapamycin
mTORC1: mechanistic target of rapamycin complex 1
mTORC2: mechanistic target of rapamycin complex
2 Myh7: myosin heavy chain 7
NaCl: sodium chloride NOS2 nitric oxide synthase 2
PBS: phosphate-buffered saline PEG polyethylene glycol
pAKT: phosphorylated
AKT: pP70S6K phosphorylated P70S6K
P70S6K: ribosomal protein S6 kinase beta-1
qPCR: quantitative polymerase chain reaction Rapa rapamycin
Retnla: resistin-like alpha
RPMI: Roswell Park Memorial Institute medium
SEM: standard error of the mean
TBS: tris-buffered saline
TNF-α: tumor necrosis factor alpha
TSC2: tuberous sclerosis complex 2

